# selfRL: Two-Level Self-Supervised Transformer Representation Learning for Link Prediction of Heterogeneous Biomedical Networks

**DOI:** 10.1101/2020.10.20.347153

**Authors:** Xiaoqi Wang, Yaning Yang, Xiangke Liao, Lenli Li, Fei Li, Shaoliang Peng

**Affiliations:** College of Computer Science and Electronic Engineering, Hunan University, Changsha 410082, China; Computer Network Information Center, Chinese Academy of Sciences, Beijing 100850, China; School of Computer Science, National University of Defense Technology, Changsha 410073, China; Peng Cheng Lab, Shenzhen 518000, China

## Abstract

Predicting potential links in heterogeneous biomedical networks (HBNs) can greatly benefit various important biomedical problem. However, the self-supervised representation learning for link prediction in HBNs has been slightly explored in previous researches. Therefore, this study proposes a two-level self-supervised representation learning, namely selfRL, for link prediction in heterogeneous biomedical networks. The meta path detection-based self-supervised learning task is proposed to learn representation vectors that can capture the global-level structure and semantic feature in HBNs. The vertex entity mask-based self-supervised learning mechanism is designed to enhance local association of vertices. Finally, the representations from two tasks are concatenated to generate high-quality representation vectors. The results of link prediction on six datasets show selfRL outperforms 25 state-of-the-art methods. In particular, selfRL reveals great performance with results close to 1 in terms of AUC and AUPR on the NeoDTI-net dataset. In addition, the PubMed publications demonstrate that nine out of ten drugs screened by selfRL can inhibit the cytokine storm in COVID-19 patients. In summary, selfRL provides a general framework that develops self-supervised learning tasks with unlabeled data to obtain promising representations for improving link prediction.

## Introduction

In recent decades, networks have been widely used to represent biomedical entities (as nodes) and their relations (as edges). Predicting potential links in heterogeneous biomedical networks (HBNs) can be beneficial to various significant biology and medicine problems, such as target identification, drug repositioning, and adverse drug reaction predictions. For example, network-based drug repositioning methods have already offered promising insights to boost the effective treatment of COVID-19 disease (Zeng et al. 2020; Xiaoqi et al. 2020), since it outbreak in December of 2019. Many network-based learning approaches have been developed to facilitate link prediction in HBNs. In particularly, network representation learning methods, that aim at converting high-dimensionality networks into a low-dimensional space while maximally preserve structural properties (Cui et al. 2019), have provided effective and potential paradigms for link prediction (Wang et al. 2018; Li et al. 2017).

Nevertheless, most of the network representation learning-based link prediction approaches heavily depend on a large amount of labeled data. The requirement of large-scale labeled data may not be met in many real link prediction for biomedical networks (Su et al. 2020). To address this issue, many studies have focused on developing unsupervised representation learning algorithms that use the network structure and vertex attributes to learn low-dimension vectors of nodes in networks (Yuxiao et al. 2020), such as GraRep (Cao, Lu, and Xu 2015), TADW (Cheng et al. 2015), LINE (Tang et al. 2015), and struc2vec (Ribeiro, Saverese, and Figueiredo 2017). However, these network presentation learning approaches are aimed at homogeneous network, and cannot applied directly to the HBNs. Therefore, a growth number of studies have integrated meta paths, which are able to capture topological structure feature and relevant semantic, to develop representation learning approaches for heterogeneous information networks. Dong *et al.* used meta path based random walk and then leveraged a skip-gram model to learn node representation (Dong, Chawla, and Swami 2017). Shi *et al.* proposed a fusion approach to integrate different representations based on different meta paths into a single representation (Shi et al. 2019). Ji *et al.* developed the attention-based meta path fusion for heterogeneous information network embedding (Ji, Shi, and Wang 2018). Wang *et al.* proposed a meta path-driven deep representation learning for a heterogeneous drug network (Xiaoqi et al. 2020). Unfortunately, most of the meta path-based network representation approaches focused on preserving vertex-level information by formalizing meta paths and then leveraging a word embedding model to learn node representation. Therefore, the global-level structure and semantic information among vertices in heterogeneous networks is hard to be fully modeled. In addition, these representation approaches is not specially designed for link prediction, thus resulting in learning an inexplicit representation for link prediction.

On the other hand, self-supervised learning, which is a form of unsupervised learning, has been receiving more and more attention. Self-supervised representation learning formulates some pretext tasks using only unlabeled data to learn representation vector without any manual annotations (Xiao et al. 2020). Self-supervised representation learning technologies have been widely use for various domains, such as natural language processing, computer vision, and image processing. However, very few approaches have been generalized for HBNs because the structure and semantic information of heterogeneous networks is significantly differ between domains, and the model trained on a pretext task may be unsuitable for link prediction tasks. Based on the above analysis, there are two main problems in link prediction based on network representation learning. The first one is how to design a self-supervised representation learning approach based on a great amount of unlabeled data to learn low-dimension vectors that integrate the different-view structure and semantic information of HBNs. The second one is how to ensure the pretext tasks in self-supervised representation learning be beneficial for link prediction of HBNs.

In order to overcome the mentioned issues, this study proposes a two-level self-supervised representation learning (selfRL) for link prediction in heterogeneous biomedical networks. First, a meta path detection self-supervised learning mechanism is developed to train a deep Transformer encoder for learning low-dimensional representations that capture the path-level information on HBNs. Meanwhile, selfRL integrates the vertex entity mask task to learn local association of vertices in HBNs. Finally, the representations from the entity mask and meta path detection is concatenated for generating the embedding vectors of nodes in HBNs. The results of link prediction on six datasets show that the proposed selfRL is superior to 25 state-of-the-art methods.

In summary, the contributions of the paper are listed below:

- We proposed a two-level self-supervised representation learning method for HBNs, where this study integrates the meta path detection and vertex entity mask self-supervised learning task based on a great number of unlabeled data to learn high quality representation vector of vertices.
- The meta path detection self-supervised learning task is developed to capture the global-level structure and semantic feature of HBNs. Meanwhile, vertex entity-masked model is designed to learn local association of nodes. Therefore, the representation vectors of selfRL integrate two-level structure and semantic feature of HBNs.
- The meta path detection task is specifically designed for link prediction. The experimental results indicate that selfRL outperforms 25 state-of-the-art methods on six datasets. In particular, selfRL reveals great performance with results close to 1 in terms of AUC and AUPR on the NeoDTI-net dataset.

## Related work

### Heterogeneous biomedical network

A heterogeneous biomedical network is defined as *G* = (*V, E*) where *V* denotes a biomedical entity set, and *E* represents a biomedical link set. In a heterogeneous biomedical network, using a mapping function of vertex type *ϕ*(*v*): *V* → *A* and a mapping function of relation type *ψ*(*e*): *E* → *R* to associate each vertex *v* and each edge *e*, respectively. *A* and *R* denote the sets of the entity and relation types, where |*A*| + |*R*| > 2.

### The schema of heterogeneous biomedical network

For a given heterogeneous network *G* = (*V, E*), the network schema *T_G_* can be defined as a directed graph defined over object types *A* and link types *R*, that is, *T*_*G*_ = (*A, R*). The schema of a heterogeneous biomedical network expresses all allowable relation types between different type of vertices, as shown in Figure 1.

**Figure 1:**
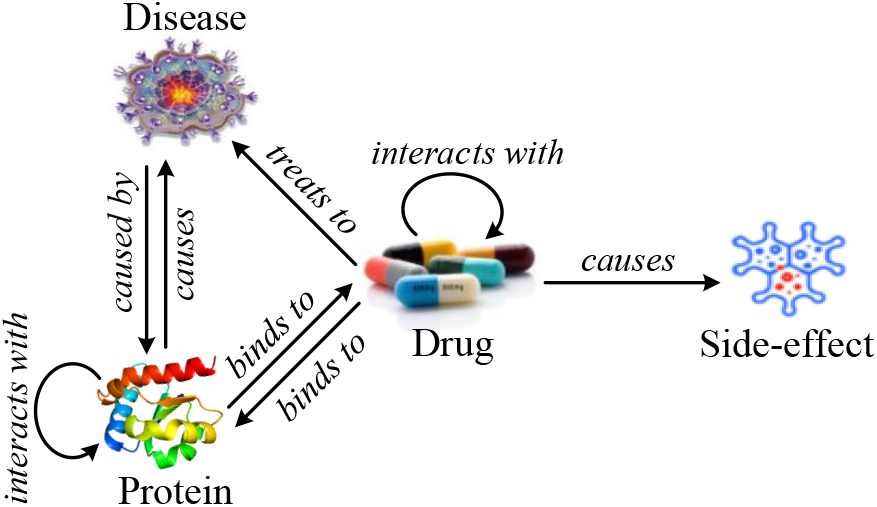
Schema of the heterogeneous biomedical network that includes four types of vertices (i.e., drug, protein, disease, and side-effect).

### Network representation learning

Network representation learning plays a significant role in various network analysis tasks, such as community detection, link prediction, and node classification. Therefore, network representation learning has been receiving more and more attention during recent decades. Network representation learning aims at learning low-dimensional representations of network vertices, such that proximities between them in the original space are preserved (Cui et al. 2019).

The network representation learning approaches can be roughly categorized into three groups: matrix factorization-based network representation learning approaches, random walk-based network representation learning approaches, and neural network-based network representation learning approaches (Yue et al. 2019). The matrix factorization-based network representation learning methods extract an adjacency matrix, and factorize it to obtain the representation vectors of vertices, such as, Laplacian eigenmaps (Belkin and Niyogi 2002) and the locally linear embedding methods (Roweis and Saul 2000). The traditional matrix factorization has many variants that often focus on factorizing the high-order data matrix, such as, GraRep (Cao, Lu, and Xu 2015) and HOPE (Ou et al. 2016). Inspired by the word2vec (Mikolov et al. 2013) model, many random walk-based representation learning models have been proposed recently, including DeepWalk (Perozzi, Alrfou, and Skiena 2014), node2vec (Grover and Leskovec 2016), and metapath2vec/metapath2vec++ (Dong, Chawla, and Swami 2017), in which a network is transformed into node sequences. These models were later extended by struc2vec (Ribeiro, Saverese, and Figueiredo 2017) for the purpose of better modeling the structural identity. Over the past years, neural network models have been widely used in various domains, and they have also been applied to the network representation learning areas. In neural network-based network representation learning, different methods adopt different learning architectures and various network information as input. For example, the LINE (Tang et al. 2015) aims at embedding by preserving both local and global network structure properties. The SDNE (Wang, Cui, and Zhu 2016) and DNGR (Cao 2016) were developed using deep autoencoder architecture. The GraphGAN (Wang et al. 2017) adopts generative adversarial networks to model the connectivity of nodes.

## Method

Predicting potential links in HBNs can greatly benefit various important biomedical problems. This study proposes selfRL that is a two-level self-supervised representation learning algorithm, to improve the quality of link prediction. The flowchart of the proposed selfRL is shown in Figure 2. Considering meta path reflecting heterogeneous characteristics and rich semantics, selfRL first uses a random walk strategy guided by meta-paths to generate node sequences that are treated as the true paths of HBNs. Meanwhile, an equal number of false paths is produced by randomly replacing some of the nodes in each of true path. Then, based on the true paths, this work proposes vertex entity masked as self-supervised learning task to train deep Transformer encoder for learning entity-level representations. In addition, a meta path detection-based self-supervised learning task based on all true and false paths is designed to train a deep Transformer encoder for learning path-level representation vectors. Finally, the representations obtained from the two-level self-supervised learning task are concatenated to generate the embedding vectors of vertices in HBNs, and then are used for link prediction.

**Figure 2:**
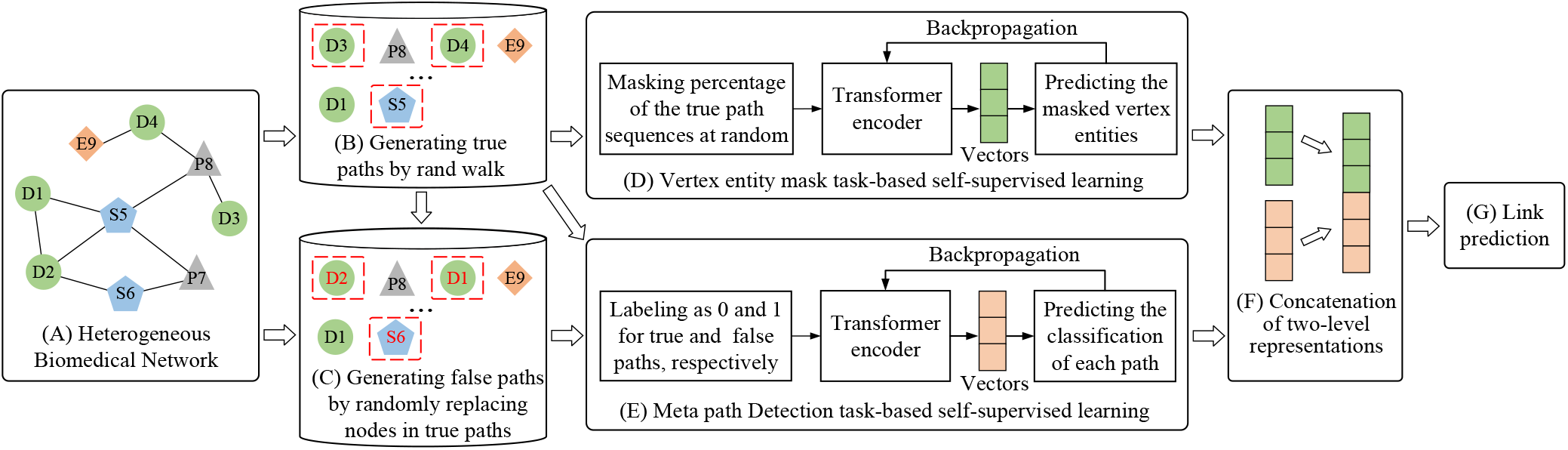
The flowchart of the proposed selfRL where the circle, triangle, rhombus, and pentagon denote the different types of node entities in the heterogeneous biomedical network.

### Meta path detection-based self-supervised learning task

#### True path generation

A meta-path is a composite relation denoting a sequence of adjacent links between nodes *A*_1_ and *A*_*i*_ in a heterogeneous network, and can be expressed in the form of 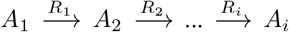, where *R*_*i*_ represents a schema between two objects. Different adjacent links indicate distinct semantics. In this study, all the meta paths are reversible, and no longer than four nodes. This is based on the results of the previous studies that meta paths longer than four nodes may be too long to contribute to the informative feature (Fu et al. 2016). In addition, Sun *et al.* have suggested that short meta paths are good enough, and that long meta paths may even reduce the quality of semantic meanings (Sun et al. 2011).

In this work, each network vertex and meta path are regarded as vocabulary and sentence, respectively. Indeed, a large percentage of meta paths are biased to highly visible objects. Therefore, three key steps are defined to keep a balance between different semantic types of meta paths, and they are as follows: (1) generate all sequences according to meta paths whose positive and reverse directional sampling probabilities are the same and equal to 0.5. (2) count the total number of meta paths of each type, and calculate their median value (*N*); (3) randomly select *N* paths if the total number of meta paths of each type is larger than *N*; otherwise, select all sequences. The selected paths are able to reflect topological structures and interaction mechanisms between vertices in HBNs, and will be used to design self-supervised learning task to learn low-dimensional representations of network vertices.

#### False path generation

The paths selected using the above procedure are treated as the true paths in HBNs. The equal number of false paths are produced by randomly replacing some nodes in each of the true paths. In other words, each true path corresponds to a false path. There is no relation between the permutation nodes and context in false paths, and the number of replaced nodes is less than the length of the current path. For instance, a true path (i.e., D3 P8 D4 E9) is shown in Figure 2(B). During the generation procedure of false paths, the 1st and 3rd tokens that correspond to D3 and D4, respectively, are randomly chosen, and two nodes from the HBNs which correspond to D2 and D1, respectively, are also randomly chosen. If there is a relationship between D2 and P8, node D3 is replaced with P2. If there is a relationship between D2 and P8, another node from the network is chosen until the mentioned conditions are satisfied. Similarly, node D4 is replaced with D1, because there are no relations between D1 and E9 (or P8). Finally, the path (i.e., D2 P8 D1 E9) is treated as a false path.

#### Meta path detection

In general language understanding evaluation, the Corpus of Linguistic Acceptability (CoLA) is a binary classification task, where the goal is to predict whether a sentence is linguistically acceptable or not (Wang et al. 2019). In addition, Perozzi *et al.* have suggested that paths generated by short random walks can be regarded as short sentences (Perozzi, Alrfou, and Skiena 2014). Inspired by their work, this study assumes that true paths can be treated as linguistically acceptable sentences, while the false paths can be regarded as linguistically unacceptable sentences. Based on this hypothesis, we proposes the meta path detection task where the goal is to predict whether a path is acceptable or not. In the proposed selfRL, a set of true and false paths is fed into the deep Transformer encoder for learning path-level representation vector. selfRL maps a path of symbol representations to the output vector of continuous representations that is fed into the softmax function to predict whether a path is a true or false path.

Apparently, the only distinction between true and false paths is whether there is an association between nodes of path sequence. Therefore, the meta path detection task is the extension of the link prediction to a certain extent. Especially, when a path includes only two nodes, the meta path detection is equal to the link prediction. For instance, judging whether a path is a true or false path, e.g., D1 S5 in Figure B, is the same as predicting whether there is a relation between D1 and S5. However, the meta path detection task is generally more difficult compared to link prediction, because it requires the understanding of long-range composite relationships between vertices of HBNs. Therefore, the meta path detection-based self-supervised learning task encourages to capturing high-level structure and semantic information in HBNs, thus facilitating the performance of link prediction.

### Vertex entity mask-based self-supervised learning task

In order to capture the local information on HBNs, this study develops the vertex entity mask-based self-supervised learning task, where nodes in true paths are randomly masked, and then predicting those masked nodes. The vertex entity mask task has been widely applied to natural language processing. However, using the vertex entity mask task to drive the heterogeneous biomedical network representation model is a less explored research. In this work, the vertex entity mask task fellows the implementation described in the BERT, and the implementation is almost identical to the original (Devlin et al. 2018). In brief, 15% of the vertex entities in true paths are randomly chosen for prediction. For each selected vertex entity, there are three different operations for improving the model generalization performance. The selected vertex entity is replaced with the ¡MASK¿ token for 80% time, and is replaced with a random node for 10% time. Furthermore, it has 10% chance to keep the original vertex.

Finally, the masked path is used for training a deep Transformer encoder model according to the vertex entity mask task where the last hidden vectors corresponding to the mask vertex entities are fed into the softmax function to predict their original vertices with cross entropy loss. The vertex entity mask task can keep a local contextual representation of every vertex.

### Two-level representations vector

The vertex entity mask-based self-supervised learning task captures the local association of the vertex in HBNs. The meta path detection-based self-supervised learning task enhances the global-level structure and semantic features of the HBNs. Therefore, the two-level representations are concatenated as the final embedding vectors that integrate structure and semantics information on HBNs from different level, as shown in Figure 2(F).

### Network architecture of selfRL

The model of selfRL is a deep Transformer encoder, and the implementation is almost identical to the original (Vaswani et al. 2017). The selfRL follows the overall architecture that includes the stacked self-attention and point-wise, fully connected layers, and softmax function, as shown in Figure 3.

**Figure 3:**
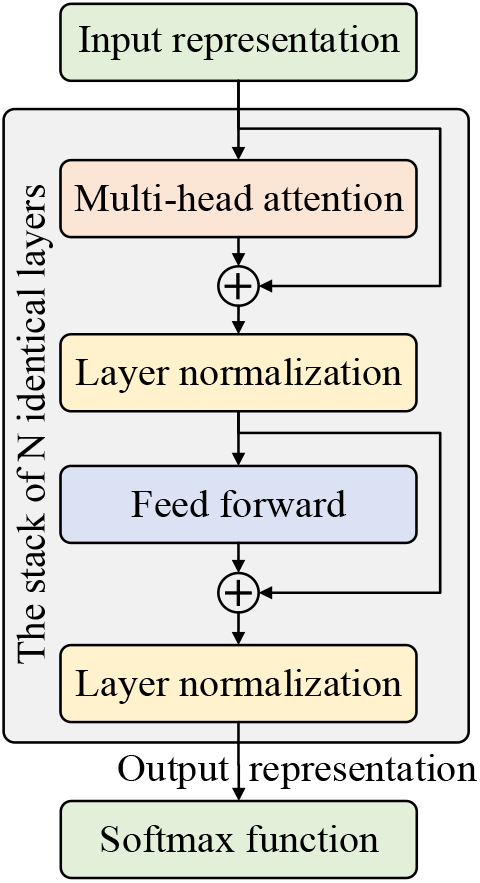
Model architecture of selfRL.

#### Multi-head attention

An attention function can be described as mapping a query vectors and a set of key-value pairs to an output vectors. The multi-head attention leads that the model can jointly attend to view from various embedding subspaces at various positions.

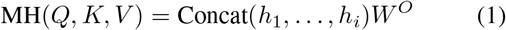

where *W*^*O*^ is a parameter matrices, and *h*_*i*_ is the attention function of *i*-th subspace, and is given as follows:

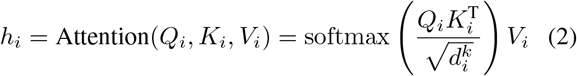

where 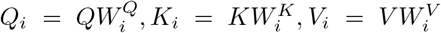 respectively denotes the query, key, and value representations of the *i*-th subspace, and 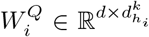, 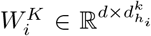 and 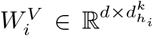 is parameter matrices which represent that *Q, K*, and *V* are transformed into *h*_*i*_ subspaces, and *d* and 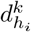 represent the dimensionality of the model and *h*_*i*_ sub-model.

#### Position-wise feed-forward network

In addition to multi-head attention layers, the proposed selfRL model include a fully connected feed-forward network, which includes two linear transformations with a ReLU activation function, is given as follows:

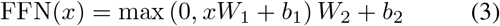

There are the same the linear transformations for various positions, while these linear transformations use various parameters from layer to layer.

#### Residual connection

For each sub-layer, a residual connection and normalization mechanism are employed. That is, the output of each sub-layer is given as follows:

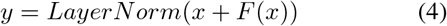

where *x* and *F* (*x*) stand for input and the transformational function of each sub-layer, respectively.

## Result

In this work, the performance of selfRL is evaluated comprehensively by link prediction on six datasets. The results of selfRL is also compared with the results of 25 methods.

### Datasets

Six datasets, including NeoDTI-Net, deepDR-Net, CTD-DDA, STRING-PPI, DrugBank-DDI, and NDFRT-DDA are used in the experiments; NeoDTI-Net and deepDR-Net were proposed in NeoDTI (Wan et al. 2018) and deepDR (Zeng et al. 2019), and the other datasets were proposed in BioNEV (Yue et al. 2019). Among these datasets, NeoDTI-Net and deepDR-Net are complex biomedical networks that are connected by multiple types of relations; the other datasets are single biomedical networks that include only one type of edge. It is worth noting that the deepDR-Net includes four types of objects (i.e., target, drug, disease, and side-effect), and four types of links (i.e., drug-drug interaction (DDI), drug-target interaction (DTI), drug-disease association (DDA), and drug-side-effect association (DSA). However, in the original deepDR database (Zeng et al. 2019), there are 12 types of vertices and 11 types of edges. The node and edge statistics of all datasets are shown in Table 1. The construction of each network can be found in NeoDTI (Wan et al. 2018), deepDR (Zeng et al. 2019), and BioNEV (Yue et al. 2019).

**Table 1:** The node and edge statistics of the datasets. Here, DDI, DTI, DSA, DDA, PDA, PPI represent the drug-drug interaction, drug-target interaction, drug-side-effect association, and drug-disease association, protein-disease association and protein-protein interaction, respectively.

| Network | Type of nodes | Type of edges | Number of nodes | Number of deges |
| --- | --- | --- | --- | --- |
| NeoDTI-Net | drug, protein, disease, side-effect | DDI, DTI, DDA, DSA, PPI, PDA | 12,015 | 1,895,445 |
| deepDR-Net | drug, protein, disease, side-effect | DDI, DTI, DDA, DDA | 16,677 | 686,298 |
| CTD-DDA | drug, disease | DDA | 12,765 | 92,813 |
| NDFRT-DDA | drug, disease | DDA | 13,545 | 56,515 |
| DrugBank-DDI | drug | DDI | 2,191 | 242,027 |
| STRING-PPI | protein | PPI | 15,131 | 359,776 |

### Baseline methods

For NeoDTI-Net datasets, the performance of selfRL is compared with those of seven state-of-the-art methods, including MSCMF (Yang et al. 2019), HNM (Lim et al. 2019), NeoDTI (Wan et al. 2018), DTINet (Luo et al. 2017), BLM-NII (Mei et al. 2013), DT-Hybrid (Alaimo et al. 2013), and NetLapRLS (Xia et al. 2010). The details on how to set the hyperparameters in above baseline approaches can be found in NeoDTI (Wan et al. 2018). For deepDR-Net datasets, the link prediction results generated by selfRL are compared with that of seven baseline algorithms, including deepDR (Zeng et al. 2019), DTINet (Luo et al. 2017), kernelized Bayesian matrix factorization (KBMF) (Gonen and Kaski 2014), support vector machine (SVM) (Cortes and Vapnik 1995), random forest (RF) (L 2001), random walk with restart (RWR) (Cao et al. 2014), and Katz (Singhblom et al. 2013). The details of the baseline approaches and hyperparameters selection can be seen in deepDR (Zeng et al. 2019). For single network datasets, selfRL is compared with 11 network representation methods, that is Laplacian (Belkin and Niyogi 2003), singular value decomposition (SVD), graph factorization (GF) (Ahmed et al. 2013), HOPE (Ou et al. 2016), Grarep (Cao, Lu, and Xu 2015), DeepWalk (Perozzi, Alrfou, and Skiena 2014), node2vec (Grover and Leskovec 2016), struc2vec (Ribeiro, Saverese, and Figueiredo 2017), LINE (Tang et al. 2015), SDNE (Wang, Cui, and Zhu 2016), and GAE (Kipf and Welling 2016). More implementation details can be found in BioNEV (Yue et al. 2019). The hyperparameters selection of baseline methods were set to default values, and the original data of NeoDTI (Wan et al. 2018), deepDR (Zeng et al. 2019), and BioNEV (Yue et al. 2019) were used in the experiments.

### Experiment settings and evaluations

The parameters of the proposed selfRL follows those of the BERT (Devlin et al. 2018) which the number of Transformer blocks (*L*), the number of self-attention heads (*A*), and the hidden size (*H*) is set to 12, 12, and 768, respectively. For the NeoDTI-Net dataset, the embedding vectors are fed into the inductive matrix completion model (IMC) (Jain and Dhillon 2013) to predict DTI. The number of negative samples that are randomly chosen from negative pairs, is ten times that of positive samples according to the guidelines in NeoDTI (Wan et al. 2018). Then, to reduce the data bias, the ten-fold cross-validation is performed repeatedly ten times, and the average value is calculated. For the deepDR-Net dataset, a collective variational autoencoder (cVAE) is used to predict DDA. All positive samples and the same number of negative samples that is randomly selected from unknown pairs are used to train and test the model according to the guidelines in deepDR (Zeng et al. 2019). Then, five-fold crossvalidation is performed repeatedly 10 times. For NeoDTI-Net and deepDR-Net datasets, the area under precision recall (AUPR) curve and the area under receiver operating characteristic (AUC) curve are adopted to evaluate the link prediction performance generated by all approaches. For other datasets, the representation vectors are fed into the Logistic Regression binary classifier for link prediction, the training set (80%) and the testing set (20%) consisted of the equal number of positive samples and negative samples that is randomly selected from all the unknown interactions according to the guidelines in BioNEV. The performance of different methods is evaluated by accuracy (ACC), AUC, and F1 score.

### Result and analysis on complex heterogeneous network

The overall performances of all methods for DTI prediction on the NeoDTI-Net dataset are presented in Figure 4. selfRL shows great results with the AUC and AUPR value close to 1, and significantly outperformed the baseline methods. In particular, NeoDTI and DTINet were specially developed for the NeoDTI-Net dataset. However, selfRL is still superior to both NeoDTI and DTINet, improving the AUPR by approximately 10% and 15%, respectively.

**Figure 4:**
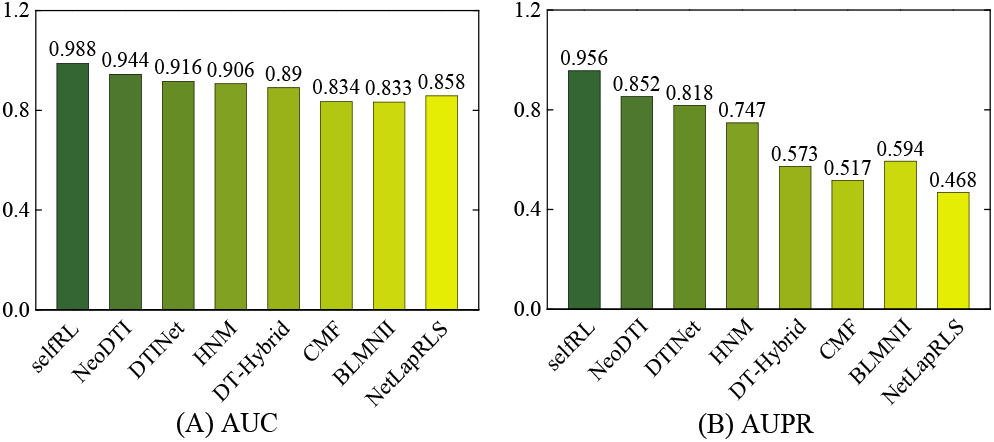
The DTI prediction results of selfRL and baseline methods for NeoDTI-Net dataset.

The results of DDA prediction of selfRL and baseline methods are represented in Figure 5. These experimental results demonstrate that selfRL generates better results of the DDA prediction on the deepDR-Net dataset than the baseline methods. However, selfRL achieves the improvements in term of AUC and AUPR less than 2%. A major reason for such a poor superiority of the selfRL to the other methods is that selfRL considers only four types of objects and edges. However, deepDR included 12 types of vertices and 11 types of edges of drug-related data. In addition, deepDR specially integrated multi-modal deep autoencoder (MDA) and cVAE model to improve the DDA prediction on the deepDR-Net dataset. Unfortunately, the selfRL+cVAE combination maybe reduce the original balance between the MDA and cVAE.

**Figure 5:**
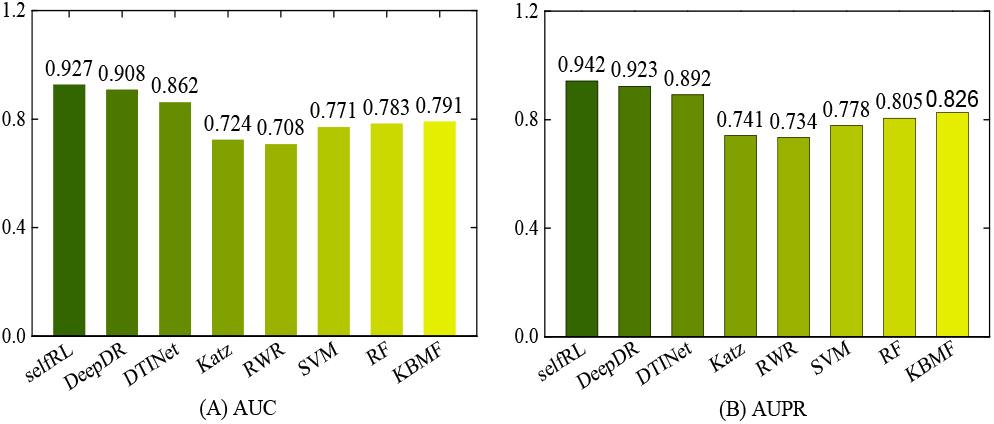
The DDA prediction results of selfRL and baseline methods for deepDR-Net dataset.

The above results and analysis indicate that the proposed selfRL is a powerful network representation approach for complex heterogeneous networks, and that can achieve very promising results in link prediction. Such a good performance of the proposed selfRL is due to the following facts: (1) selfRL designs a two-level self-supervised learning task to integrate the local association of a node and the global level information of HBNs. (2) meta path detection self-supervised learning task that is an extension of link prediction, is specially designed for link prediction. In particular, path detection of two nodes is equal to link prediction. Therefore, the representation generated by meta path detection is able to facilitate the link prediction performance. (3) selfRL uses meta paths to integrate the structural and semantic features of HBNs.

### Result and analysis on single network

In this section, the link prediction results on four single network datasets are presented to further verify the representation performance of the proposed selfRL. The link prediction results of selfRL and baseline methods on four single network datasets are listed in Table 2, and the best results are marked in **boldface**. selfRL shows higher accuracy in link prediction on four single networks compared to the other 11 baseline approaches. Especially, the proposed selfRL can achieves an approximately 2% improvement in terms of AUC and ACC over the second best method on the STRING-PPI dataset. The AUC value of link prediction on the NDFRT-DDA dataset is improved from 0.963 to 0.971 when selfRL is compared with GraRep. However, GraRep only achieves an enhancement of 0.001 compared to LINE that is the third best method on the STRING-PPI dataset. Therefore, the improvement of selfRL is significant in comparison to the enhancement of GraRep compared to LINE. Meanwhile, we also notice that selfRL have poor superiority to the second best method on the CTD-DDA and DrugBank-DDI datasets. One possible reason for this result can be that the structure and semantic of the CTD-DDA and DrugBank-DDI datasets are simple and monotonous, so most of the network representation approaches are able to achieve good performance on them. Consequently, The proposed selfRL is a potential representation method for the single network datasets, and can contribute to link prediction by introducing a two-level self-supervised learning task.

**Table 2:** Results of link prediction of selfRL and baseline methods on CTD-DDA, NDFRT-DDA, DrugBank-DDI, and STRING-PPI datasets.

| Method | CTD-DDA |  |  | NDFRT-DDA |  |  | DrugBank-DDI |  |  | STRING-PPI |  |  |
| --- | --- | --- | --- | --- | --- | --- | --- | --- | --- | --- | --- | --- |
|  | AUC | ACC | F1 | AUC | ACC | F1 | AUC | ACC | F1 | AUC | ACC | F1 |
| selfRL | **0.966** | **0.906** | **0.907** | **0.971** | **0.938** | **0.939** | **0.930** | **0.853** | **0.858** | **0.926** | **0.856** | **0.858** |
| Laplacian | 0.856 | 0.793 | 0.802 | 0.930 | 0.917 | 0.921 | 0.796 | 0.720 | 0.729 | 0.639 | 0.596 | 0.586 |
| SVD | 0.936 | 0.855 | 0.854 | 0.779 | 0.707 | 0.700 | 0.919 | 0.837 | 0.837 | 0.867 | 0.794 | 0.790 |
| GF | 0.884 | 0.808 | 0.805 | 0.720 | 0.660 | 0.655 | 0.882 | 0.720 | 0.802 | 0.810 | 0.746 | 0.747 |
| HOPE | 0.951 | 0.886 | 0.887 | 0.949 | 0.928 | 0.931 | 0.923 | 0.844 | 0.846 | 0.839 | 0.764 | 0.764 |
| GraRep | 0.960 | 0.899 | 0.900 | 0.963 | 0.931 | 0.934 | 0.925 | 0.845 | 0.846 | 0.894 | 0.823 | 0.822 |
| DeepWalk | 0.929 | 0.866 | 0.864 | 0.783 | 0.710 | 0.709 | 0.921 | 0.840 | 0.842 | 0.884 | 0.813 | 0.814 |
| node2vec | 0.911 | 0.838 | 0.835 | 0.819 | 0.742 | 0.741 | 0.902 | 0.819 | 0.819 | 0.828 | 0.758 | 0.756 |
| struc2vec | 0.965 | 0.903 | 0.903 | 0.958 | 0.913 | 0.921 | 0.904 | 0.826 | 0.830 | 0.909 | 0.838 | 0.841 |
| LINE | 0.965 | 0.904 | 0.904 | 0.962 | 0.934 | 0.935 | 0.905 | 0.825 | 0.829 | 0.859 | 0.788 | 0.795 |
| SDNE | 0.935 | 0.863 | 0.861 | 0.944 | 0.896 | 0.897 | 0.911 | 0.833 | 0.838 | 0.884 | 0.813 | 0.814 |
| GAE | 0.937 | 0.857 | 0.856 | 0.813 | 0.735 | 0.730 | 0.917 | 0.836 | 0.840 | 0.900 | 0.827 | 0.829 |

### Performance of selfRL by ablation analysis

In the NeoDTI and deepDR, low-dimensional representations of nodes in HBNs are first learned by network representation approaches, and then are fed into classifier models for predicting potential link among vertices. To further examine the contribution of the network representation approaches, the low-dimensional representation vector is fed into SVM that is a traditional and popular classifier for link prediction. The experimental results of these combinations are shown in Table 3. selfRL achieves the best performance in link prediction for complex heterogeneous networks, providing a great improvement of over 10% with regard to AUC and AUPR compared to the NeoDTI and deepDR. With the change of classifiers, the result of selfRL in link prediction reduced from 0.988 to 0.962 on the NeoDTI-Net dataset, while the AUC value of NeoDTI approximately reduce by 9%. Interestingly, The results on the deepDR-Net dataset are similar. Therefore, the experimental results indicate that the network representation performance of selfRL is more robust and better than those of the other embedding approaches. This is mainly because selfRL integrates a two-level self-supervised learning model to fuse the rich structure and semantic information from different views. Meanwhile, path detection is an extension of link prediction, yielding to better representation in link prediction.

**Table 3:** Performances of selfRL and baseline methods compared with the traditional SVM classifier on NeoDTI-Net and DeepDR-Net datasets, respectively.

| Method | NeoDTI-Net |  | Method | DeepDR-Net |  |
| --- | --- | --- | --- | --- | --- |
|  | AUC | AUPR |  | AUC | AUPR |
| selfRL | **0.962** | **0.948** | selfRL | **0.887** | **0.899** |
| NeoDTI | 0.855 | 0.883 | DeepDR | 0.773 | 0.779 |

### selfRL-based drug repositioning for anti-inflammatory response in COVID-19

The emergence and rapid expansion of COVID-19 have posed a global health threat. Recent studies have demonstrated that the cytokine storm, namely the excessive inflammatory response, is a key factor leading to death in patients with COVID-19. Therefore, it is urgent and important to discover potential drugs that prevent the cytokine storm in COVID-19 patients. Meanwhile, it has been proven that interleukin(IL)-6 is a potential target of anti-inflammatory response, and drugs targeting IL-6 are promising agents blocking cytokine storm for severe COVID-19 patients (Mehta et al. 2020). In the experiments, selfRL is used for drug repositioning for COVID-19 disease which aim to discovery agents binding to IL-6 for blocking cytokine storm in patients. The low-dimensional representation vectors generated by selfRL are fed into the IMC algorithm for predicting the confidence scores between IL-6 and each drug in NeoDTI-Net dataset. Then, the top-10 agents with the highest confidence scores are selected as potential therapeutic agents for COVID-19 patients. The 10 candidate drugs and their anti-inflammatory mechanisms of action *in silico* is shown in Table 4.

**Table 4:** Ten high-confidence drugs and their anti-inflammatory mechanisms of action *in silico*. NA denotes that the anti-inflammatory ability of agents has not been found in PubMed publications.

| No. | Drugbank ID | Drug Name | Anti-inflammatory mechanism of action | PMID |
| --- | --- | --- | --- | --- |
| 1 | DB01254 | Dasatinib | Reducing the mRNA levels | 19786067 |
| 2 | DB00619 | Imatinib | Inhibiting the function of human monocytes | 21938724, 15661041 |
| 3 | DB01169 | Arsenic trioxide | Unclear | 22465848, 16638192 |
| 4 | DB01136 | Carvedilol | Reducing the mRNA levels | 27746314, 12048901, 17420794 |
| 5 | DB01029 | Irbesartan | Unclear | 15655130, 15867169 |
| 6 | DB00539 | Toremifene | NA | NA |
| 7 | DB00594 | Amiloride | Unclear | 8770057 |
| 8 | DB00571 | Propranolol | Unclear | 31849395, 31542162, 28114591 |
| 9 | DB00328 | Indomethacin | Reducing the mRNA levels | 1611085, 14991990, 32412158 |
| 10 | DB00398 | Sorafenib | Unclear | 31387809, 30154433 |

The knowledge from PubMed publications demonstrates that nine out of ten drugs are able to reduce the release and express of IL-6 for exerting anti-inflammatory effects *in silico*. Meanwhile, there are three drugs (i.e., dasatinib, carvedilol, and indomethacin) that inhibit the release of IL-6 by reducing the mRNA levels of IL-6. However, imatinib inhibits the function of human monocytes to prevent the expression of IL-6. In addition, although the anti-inflammatory mechanisms of action of five agents (i.e., arsenic trioxide, irbesartan, amiloride, propranolol, sorafenib) are uncertain, these agents can still reduce the release or expression of IL-6 for preforming anti-inflammatory effects. Therefore, the top ten agents predicted by selfRL-based drug repositioning is able to be used for inhibiting cytokine storm in patients with COVID-19, and should be taken into consideration in clinical studies on COVID-19. These results further indicate that the proposed selfRL is a powerful network representation learning approach, and can facilitate the link prediction in HBNs.

### Different level representation performance analysis of selfRL

In this study, selfRL uses Transformer encoders to learn representation vectors by the proposed vertex entity mask and meta path detection tasks. Meanwhile, the entity- and path-level representations are concatenated for generating the embedding vector of nodes in HBNs. In order to demonstrate the contribution of the different level representation, this work compared selfRL with several representation combinations of the last hidden (LH) or last two hidden (LTH) layers of the trained Transformer. The link prediction result of different level representations are shown in Table 5. selfRL achieves the best performance. Meanwhile, the results show that the two-level representation are superior to the single level representation. Interestingly, the concatenation of vectors from the LTH layers is beneficial to improving the link prediction performance compared to the mean value of the vectors from the LTH layers for each level representation model. This is intuitive since two-level representation can fuse the structural and semantic information from different views in HBNs. Meanwhile, larger number of dimensions can provide more and richer information.

**Table 5:** The DTI and DDA prediction result of selfRL and baseline methods on the NeoDTI-Net and DeepDR-Net datasets. The MLTH and CLTH stand for the mean and concatenation values of representation from the last two hidden layers, respectively. ATLRe denotes the mean value of the two-level representation from the last hidden layer.

| Different level vector |  | NeoDTI-Net |  | DeepDR-Net |  |
| --- | --- | --- | --- | --- | --- |
|  |  | AUC | AUPR | AUC | AUPR |
| entity-level | LH | 0.911 | 0.898 | 0.774 | 0.785 |
|  | ALTH | 0.913 | 0.902 | 0.796 | 0.808 |
|  | CLTH | 0.916 | 0.907 | 0.797 | 0.809 |
| path-level | LH | 0.832 | 0.822 | 0.781 | 0.781 |
|  | ALTH | 0.828 | 0.810 | 0.802 | 0.805 |
|  | CLTH | 0.866 | 0.853 | 0.804 | 0.809 |
| Two-level | ATLRe | 0.956 | 0.941 | 0.872 | 0.875 |
|  | selfRL | **0.962** | **0.948** | **0.887** | **0.899** |

## Conclusion

This study proposes a two-level self-supervised representation learning, termed selfRL, for link prediction in heterogeneous biomedical networks. The proposed selfRL designs a meta path detection-based self-supervised learning task, and integrates vertices entity-level mask tasks to capture the rich structure and semantics from two-level views of HBNs. The results of link prediction indicate that selfRL is superior to 25 state-of-the-art approaches on six datasets. In the future, we will design more self-supervised learning tasks with unable data to improve the representation performance of the model. In addition, we will also developed the effective multi-task learning framework in the proposed model.

## Acknowledgments

This work was supported by National Key R&D Program of China 2017YFB0202602, 2018YFC0910405, 2017YFC1311003, 2016YFC1302500, 2016YFB0200400, 2017YFB0202104; NSFC Grants 81973244, U19A2067, 61772543, U1435222, 61625202, 61272056; Science Foundation for Distinguished Young Scholars of Human Province (2020JJ2009); The Funds of Peng Cheng Lab, State Key Laboratory of Chemo/Biosensing and Chemometrics; the Fundamental Research Funds for the Central Universities, and Guangdong Provincial Department of Science and Technology under grant No. 2016B090918122.

